# Knockdown of the E3 Ubiquitin ligase UBR5 and its role in skeletal muscle anabolism

**DOI:** 10.1101/2020.06.05.116145

**Authors:** Daniel C. Turner, David C. Hughes, Leslie M. Baehr, Robert A. Seaborne, Mark Viggars, Jonathan C. Jarvis, Piotr P. Gorski, Claire E. Stewart, Daniel J. Owens, Sue C. Bodine, Adam P. Sharples

**Author notes:** Corresponding Authors Details: Sue C. Bodine & Adam P. Sharples, &. Primary Authors. These authors contributed equally to this work.

## Abstract

UBR5 is an E3-ubiquitin-ligase positively associated with anabolism, hypertrophy and recovery from atrophy in skeletal muscle. The precise mechanisms underpinning UBR5’s role in the regulation of skeletal muscle mass remain unknown. The present study aimed to elucidate these mechanisms by silencing the UBR5 gene *in-vitro* and *in-vivo*. The siRNA-induced reduction (−77%) in UBR5 gene expression in human myotubes was prevented by mechanical loading, suggesting that UBR5 gene expression may be regulated via mechano-transduction signalling. Therefore, we electroporated a UBR5-RNAi plasmid into mouse tibialis anterior muscle *in-vivo* to investigate the impact of reduced UBR5 on mechano-transduction signalling MEK/ERK/p90RSK and Akt/p70S6K/4E-BP1/rpS6 pathways. Seven days post UBR5 RNAi electroporation, while reductions in overall muscle mass were not detected, mean CSA of GFP-positive fibers was reduced (−9.5%) and the number of large fibers was lower versus the control. Importantly, UBR5-RNAi significantly reduced total RNA, muscle protein synthesis, ERK1/2 and Akt phosphorylation. Whilst p90RSK phosphorylation significantly increased, total p90RSK protein levels demonstrated a 45% reduction with UBR5-RNAi. Finally, these early signalling events after 7 days of UBR5 knockdown culminated in significant reductions in muscle mass (−4.6%) and larger reductions in fiber CSA (−18.5%) after 30 days. This was associated with increased levels of the phosphatase, PP2Ac, and inappropriate chronic elevation of p70S6K and rpS6 between 7 and 30 days, and corresponding reductions in eIF4e. This study demonstrates UBR5 plays an important role in anabolism/hypertrophy, whereby knockdown of UBR5 culminates in skeletal muscle atrophy.

## Introduction

The regulation of skeletal muscle (SkM) mass is orchestrated by the activity of key signalling pathways that control protein breakdown and synthesis within myofibers. The breakdown or atrophy of SkM mass is mediated, in-part, by the ubiquitin-proteasome system (3, 7), which is composed of three key enzymes that activate and conjugate (E1 & E2 enzymes) small ubiquitin molecules to target protein substrates (E3 ligases) for recognition and subsequent degradation in the 26S proteasome. The most characterised E3 ubiquitin ligases associated with SkM atrophy are the muscle specific RING finger protein 1 (MuRF1 or Trim63) and the F-box containing ubiquitin protein ligase atrogin-1 (Atrogin-1; Gomes *et al.*, 2001), otherwise known as muscle atrophy F-box (MAFbx; Bodine *et al.*, 2001*a*).

Interestingly, recent work identified that a HECT domain E3 ligase named, ubiquitin protein ligase E3 component n-recognin 5 (EDD1 or UBR5) was significantly altered at the DNA methylation and gene expression level following resistance exercise (RE) in human SkM (36, 37). UBR5 DNA methylation decreased (hypomethylated) and mRNA expression increased after 7 weeks of training-induced SkM hypertrophy, with further enhanced changes reported following a later 7 weeks of retraining (36, 37). The pattern observed in DNA methylation and gene expression, which significantly correlated with changes in lean mass, suggested that there may be a role for UBR5 during muscle hypertrophy in contrast to E3 ligases, MuRF1 and MAFbx that are associated with atrophy. Additional work has provided further support for this hypothesis, whereby UBR5 expression significantly increased after acute mechanical loading in bioengineered SkM *in-vitro*, in response to synergistic ablation/functional overload (FO), and after programmed resistance training in rodent muscle *in-vivo*, yet with no change in MuRF1 and MAFbx expression (35). UBR5 also increased during recovery from hindlimb unloading (HU) and tetrodotoxin induced-disuse atrophy, again with no increase in MuRF1 and MAFbx (35). Furthermore, increased gene expression of UBR5 in these models resulted in greater abundance of UBR5 protein content following FO-induced hypertrophy of the mouse plantaris muscle *in-vivo*, and over the time-course of regeneration in primary human muscle cells *in-vitro* (35). A recent study also supported the role of UBR5 as being essential for muscle growth through RNAi screening in *Drosophila* larvae, where UBR5 inhibition led to smaller sized larvae (23). Collectively, these data support the notion that UBR5 is involved in load-induced anabolism and hypertrophy, in contrast with the well-known MuRF1 and MAFbx E3 ligases that are associated with muscle atrophy.

The most characterised signalling pathway involved in anabolism, protein synthesis and hypertrophy of SkM is the Akt/mTORC1/p70S6K pathway (Baar *et al.*, 2000; Bodine *et al.*, 2001; Goodman *et al.*, 2011). Further work by the Esser laboratory also reported PI3K/Akt-independent activation of mTORC via mitogen-activated protein kinase (MEK)/extracellular signal-regulated kinase (ERK) signalling which was critical for overload-induced hypertrophy (28). Earlier work also demonstrated increased ERK1/2 phosphorylation after acute resistance exercise (RE) in humans (13) and after mechanical loading in C_2_C_12_ myotubes (21). Downstream of ERK1/2, phosphorylation of the ribosomal S6 kinase (p90RSK) has been shown to increase with acute endurance and resistance exercise in rodent and human SkM (29, 43, 45). Interestingly, previous work also suggests that UBR5 is regulated via ERK/p90RSK signalling in non-muscle cells (9). Indeed, UBR5 has been shown to be a target substrate for ERK2 in the COS-1 kidney fibroblast cell line after treatment with epidermal growth factor (EGF), an established ligand that initiates downstream ERK signalling when bound to its receptor (EGFR) (15). Others have also shown that p90RSK, phosphorylates UBR5 in HeLa cancer cells at various sites, and may therefore be involved in growth of cancer cells (9). Further, UBR5 has been suggested to target protein phosphatase 2A subunit C (PP2Ac; catalytic subunit) for proteasomal degradation which is reported to negatively regulate Akt and ERK cascades (25–27, 38). Therefore, given that ERK/p90RSK signalling is associated with load-induced SkM hypertrophy (28) and UBR5 is associated with ERK/p90RSK signalling in non-muscle cells (9, 15), assessing ERK/p90S6K signalling as well as established load-induced Akt/ p70S6K signalling, together with negative regulator PP2Ac, after experimental manipulation of UBR5 is required to elucidate UBR5s mechanistic role in positively regulating muscle mass.

Therefore, in the present study, we first aimed to silence UBR5 (via siRNA) in human myotubes during mechanical loading to understand UBR5s role in response to loading. We demonstrated that siRNA-induced silencing of UBR5 in non-loaded control conditions reduced its gene expression by ∼77%, and that this reduction in expression was completely prevented by the addition of mechanical loading. Providing positive evidence towards our hypothesis that UBR5 gene expression was perhaps regulated via mechano-transduction signalling. Therefore, to test this hypothesis we used a miR-based RNAi for UBR5, using pcDNA TM 6.2-GW/EmGFP-miR electroporated into murine tibialis anterior (TA) muscle, to investigate the impact of reduced UBR5 on MEK/ERK/p90RSK/MSK1 and PP2Ac/Akt/GSK3β/p70s6K/rpS6/eIF4e signalling *in-vivo*. UBR5 silencing led to a reduction in the CSA of transfected fibers, the frequency of larger fibers (≥3400 µm^2^), total RNA content, protein synthesis, ERK1/2 and Akt activity and total p90RSK levels. However, activity of p90RSK, p70S6K and rpS6 activity increased. Finally, the changes at 7 days also culminated in even larger reductions in muscle mass and fiber CSA after 30 days post transfection of UBR5 RNAi. This was also associated with increased PP2Ac and chronic elevation of p70S6K and rpS6 between 7 and 30 days, as well as reductions in eIF4e. Overall, the present study supports the notion that UBR5 plays an important role in muscle anabolism/hypertrophy, and that a reduction in UBR5 is associated with early reductions in ERK1/2, Akt activity, total p90RSK and protein synthesis that culminates in later atrophy via increased PP2Ac and inappropriate activation of p70S6K and rpS6 signaling.

## Methods

### Skeletal Muscle Tissue and Ethical Approval

Prior to collection of human skeletal muscle (SkM) tissue, participants gave written informed consent. Ethical approval was granted by the NHS West Midlands Black Country, UK, Research Ethics Committee (NREC approval no. 16/WM/0103). Muscle tissue was handled and stored in accordance with Human Tissue Act (2004) regulations. For *in-vitro* myotube experiments, human SkM biopsies were obtained from the vastus lateralis muscle of young healthy males (23 ± 3.2 yrs, 75.4 ± 3.7 kg, 180.7 ± 2.1 cm, BMI 23.1 ± 1.5 kg/m^2^) using a needle biopsy instrument (CR Bard, Crawley, UK) as described fully elsewhere (Turner *et al.*, 2019*a*). For *in-vivo* muscle tissue experiments, C57Bl/6 male mice between twelve and sixteen weeks-old (*n* = 10) were obtained from Charles River Laboratories for electroporation experiments. All animal procedures were approved by the Institutional Animal Care and Use Committee at the University of Iowa. During tissue collection, animals were anaesthetized with 2–3% inhaled isoflurane. On completion of tissue removal, mice were euthanised by exsanguination.

### Isolation and Culture of Human Skeletal Muscle Derived Cells (HMDC)

Following human biopsy procedures, muscle tissue was placed in sterile microfuge tubes (Ambion^®^, Thermo Fisher Scientific Denmark) containing 1.5 ml of chilled (4°C) transfer media composed of Ham’s F-10 medium including 1 mM L-glutamine (LG; Thermo Fisher Scientific, Denmark), 0.1% heat inactivated fetal bovine serum (hiFBS; Gibco™, South America Origin, Fisher Scientific, UK), 0.1% heat inactivated new born calf serum (hiNBCS; Gibco, New Zealand Origin, Fisher Scientific, UK), 100 U/ml penicillin (Lonza, UK), 100 μg/ml streptomycin (Lonza, UK), 2.5 μg/ml amphotericin B (Sigma-Aldrich, UK) and transported on ice to a class II biological safety cabinet (Kojair Biowizard Silverline, Finland) to undergo subsequent cell isolations as described in previous work by our group (10, 32, 40). Briefly, muscle tissue was minced in 5 ml trypsin (0.05%)/EDTA (0.02%) solution using 2 × sterile scalpels (No. 11, Swann-Morton, UK). The contents were transferred to a sterile specimen pot and magnetically stirred at 37°C for 10 mins. This process was repeated, and then homogenised tissue was placed in a sterile 15 ml tube (Falcon^®^, Fisher Scientific, UK) containing 2 ml heat inactivated horse serum (hiHS; Gibco™, New Zealand Origin, Fisher Scientific, UK) to neutralise the trypsin. The tube was centrifuged (340 × g for 5 min at 24°C) and the supernatant was seeded on to 2 separate pre-gelatinised (0.2% in dH_2_O; Type A, Sigma-Aldrich, UK) T25 flasks (Nunc™, Thermo Fisher Scientific, Denmark) containing 7.5 ml of growth medium (GM, composed of Ham’s F-10 medium that included 1 mM LG, 10% hiFBS, 10% hiNBCS, 4 mM LG, 100 U/ml penicillin, 100 μg/ml streptomycin and 2.5 μg/ml amphotericin B). All flasks were then incubated (HERAcell 150i CO_2_ humidified incubator, Thermo Fisher Scientific, Denmark) at 37°C and 5% CO_2_ until ∼80% confluency was attained. Cells were then transferred to T75 flasks and passaged (P7-9) until sufficient cell quantities were acquired for experimentation.

### Immunocytochemistry for Desmin Positivity of HDMC

HMDC were washed 3 × TBS (1×; Sigma-Aldrich, UK) and fixed in ice-cold methanol:acetone:TBS (25:25:50 for 15 mins) then in methanol:acetone (50:50) for a further 15 mins. Following fixation, HMDC were permeabilised in 0.2% Triton X-100 (Sigma-Aldrich, UK) and blocked in 5% goat serum (Sigma-Aldrich, UK) in TBS for 90 mins. Cells were washed 3 × in TBS and incubated overnight (4°C) in 300 μl of anti-desmin (Abcam Cat# ab15200, RRID: AB_301744) primary antibody made up in TBS, 2% goat serum and 0.2% Triton X-100 at concentrations of 1:50. After overnight incubation, the primary antibody was removed and HMDC were washed 3 × in TBS. Cells were incubated at RT for 3 hrs in 300 μl secondary antibody solution containing anti-rabbit TRITC (Sigma-Aldrich Cat# T6778, RRID: AB_261740) at a concentration of 1:75 in 1× TBS, 2% goat serum and 0.2% Triton X-100 to counterstain myoblasts. After a further 3 × TBS washes, 300 μl of DAPI (Thermo Fisher Scientific Cat# D1306, RRID: AB_2629482) was added to the cells at a concentration of 300 nM for 30 mins to counterstain myonuclei. Immunostained HMDC were visualised and imaged using confocal microscopy (Olympus IX83, Japan) with TRITC (Excitation: 557 nm, Emission: 576 nm) and DAPI (Excitation: 358 nm, Emission: 461 nm) filter cubes. Immuno-images were imported to Fiji/ImageJ (version 2.0.0) software to determine desmin positivity via counting the total number of myoblasts overlapping nuclei divided by the total number of nuclei present. Desmin positivity was the same across all participants (72 ± 2.4%).

### Transfection of UBR5 siRNA into Human Myotubes

After serial passaging, HMDC (passage 7-9) were seeded onto plastic 6-well fibronectin-coated (10 μg/ml in PBS, Sigma-Aldrich, UK) culture plates (described below) at a density of 9 × 10^4^ cells/ml in 2 ml GM. Once 80% confluency was attained, medium was switched to 2 ml differentiation media (DM; same media components as GM with the exception of lower 2% hiFBS) for the remaining 10 days. Each well received an additional 1 ml of DM at 72 hrs and 7 days. At day 10, existing media was removed and the resulting myotubes received either 750 µl of fresh DM alone or 750 µl DM containing 9 µl HiPerFectTransfection™ (Qiagen, UK) and 20 nmol Flexitube GeneSolutions™ (siUBR5, Qiagen, UK). The Flexitube GeneSolutions includes 4 × siRNA’s that target multiple regions of the 10900 bp UBR5/EDD1 human gene (see figure in results) and evoked a 77 ± 1.28% (mean ± SEM) reduction in UBR5 gene expression after 3 hours when using a 20 nmol concentration. All siRNA target sequences and locations are displayed in the results figures. Also, using a scramble siRNA (Qiagen, AllStars Negative Control siRNA™ including HiPerFectTransfection reagent) we confirm that we saw no significant reduction in UBR5 at this 3h time point. Please note that we first established the recommended 10 nmol siRNA concentration did not reduce UBR5 gene expression substantially (reduction of only 25.72 ± 4.72%) in human myotubes and that higher concentrations were therefore required during loading experiments.

### Mechanical Loading of Myotubes

HMDC were first differentiated on fibronectin-coated (10 μg/ml in PBS, Sigma-Aldrich, UK) flexible-bottomed culture plates (25 mm BioFlex^®^, Dunn Labortecknik, Germany) for 10 days as described above. Resultant myotubes were dosed with either 20 nmol UBR5 siRNA or DM alone, immediately after which, loading experiments commenced. Myotubes were subjected to equibiaxial tension using the Flexcell^®^ FX-5000™ Tension system (Dunn Labortecknik, Germany) and a low frequency (0.15 Hz with sine wave) intermittent loading regime was applied. The loading regime consisted of 4 sets × 10 repetitions with 90 s rest between sets, representing 1 loading session. This was repeated 5 times with each loading session interspersed with a 3.5 min rest, totalling a regime of 60 mins. Non-loaded control myotubes were also assembled to the Flexcell^®^ base plate, however, the vacuum entry was sealed to avoid any unwanted loading. Following cessation of mechanical loading, both loaded and non-loaded myotubes were lysed for RNA at 3 hrs. Due to the low number of human cells available for these experiments, protein extraction for signalling analysis was not possible.

### RNA Extraction, Primer Design and Polymerase Chain Reaction (PCR) (Human myotubes)

After incubation, existing media was removed and myotubes were washed 2 × in PBS before lysing cells with 300 μl TRIzol (Invitrogen™, Thermo Fisher Scientific, Denmark) for subsequent RNA extraction. RNA concentrations and purities (i.e. A_260_/A_280_ ratios) were quantified using a spectrophotometer (NanoDrop™ 2000, Thermo Fisher Scientific, Denmark). A_260_/A_280_ ratios demonstrated high quality RNA across all samples (1.91 ± 0.22, mean ± SD). A one-step PCR kit (QuantiFast™ SYBR^®^ Green, Qiagen, UK) was used to assess gene expression. Firstly, samples were diluted in nuclease-free H_2_O to ensure a concentration of 35 ng RNA in 10 μl reactions, made up of: 4.75 μl (7.37 ng/μl) RNA sample and 5.25 μl of master mix (MM) composed of 5 μl SYBR green mix, 0.1 μl of reverse transcriptase (RT) mix and 0.075 μl of both forward and reverse primers (both at 100 µM). Primers were designed using both Clustal Omega (https://www.ebi.ac.uk/Tools/msa/clustalo/) and Primer-BLAST (NCBI, https://www.ncbi.nlm.nih.gov/tools/primer-blast/) and were purchased from Sigma-Aldrich. All primer sequence information is described in Table 1. PCR reactions were transferred to a qRT-PCR thermocycler (Rotorgene 3000Q, Qiagen, UK) with supporting software (Hercules, CA, USA) to undergo amplification as follows: 10 min hold at 50°C (reverse transcription/cDNA synthesis), 95°C for 5 min (transcriptase inactivation and initial denaturation step) and PCR Steps of 40 cycles; 95°C for 10 s (denaturation), 60°C for 30 s (annealing and extension). Upon completion, melt curve analyses was carried out to identify non-specific amplification or primer-dimer issues for the reference genes (B2M and RPL13a) and UBR5. These analyses demonstrated a single melt curve/peak for all genes. Gene expression was then quantified using the DDCT (2-^ΔΔ^CT) equation (33) against the mean of 2 reference genes (B2M and RPL13a, 15.33 ± 0.7 with low variation across all conditions of 4.56%) and the mean C_T_ value derived from the non-treated DM group for each time point (0/3 hrs) and condition (non-loaded/loaded). The PCR efficiencies were similar for the reference genes, RPL13a (97.4 ± 7.75%, with 7.96% variation) and B2M (92.08 ± 3.18%, with 3.46% variation), and the gene of interest, UBR5 (90.93 ± 4.94%, with 5.43% variation).

**Table 1.**
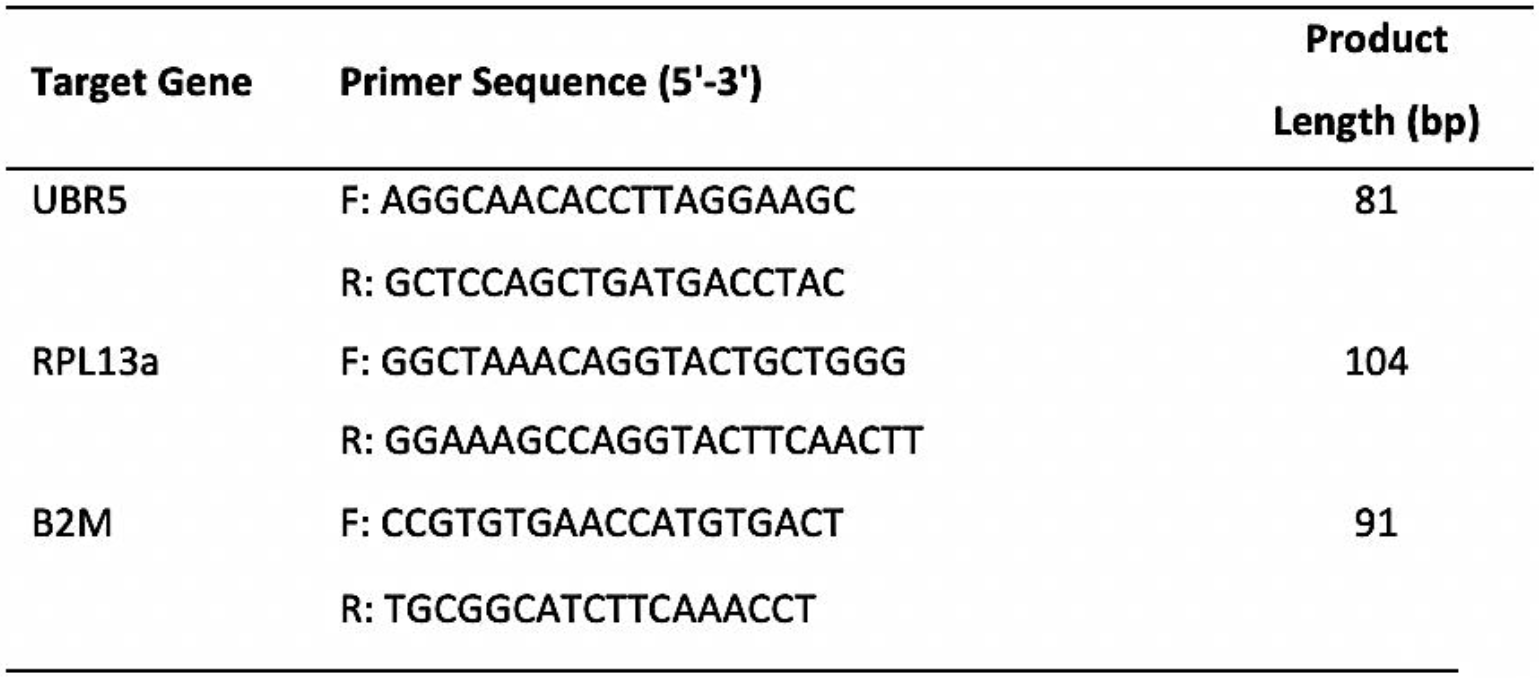
Primer sequence information for genes analysed in human myotubes.

### Plasmid Design and Electroporation of Tissue In-Vivo

An RNAi system which was designed and purchased from Invitrogen (Thermo Fisher Scientific, *USA*) was implemented. Specifically, the negative control/empty vector (EV) RNAi plasmid was described previously (14, 22, 35) and encodes emerald green fluorescent protein (EmGFP) and a non-targeting pre-miRNA under bicistronic control of the CMV promoter in the pcDNA6.2GW/EmGFP-miR plasmid (Invitrogen). The UBR5 RNAi plasmid also encodes EmGFP plus an artificial pre-miRNA targeting mouse UBR5 under bicistronic control of the CMV promoter. It was generated by ligating the Mmi571982 oligonucleotide duplex (Invitrogen) into the pcDNA6.2GW/EmGFP-miR plasmid. The UBR5 RNAi plasmid is designed to target the nucleotide sequence in the HECT domain between amino acids positions 2695-2754 (location/schematic of the RNAi on UBR5 protein structure is also included in the results). The electroporation technique was performed as previously described (35). Briefly, after a 2hr pre-treatment with 0.4 units/ul of hyaluronidase, 20 μg plasmid DNA was injected into the tibialis anterior (TA) muscle and the hind limbs were placed between two-paddle electrodes and subjected to 10 pulses (20 msec) of 175 V/cm using an ECM-830 electroporator (BTX Harvard Apparatus). Mice were injected with the UBR5 RNAi plasmid and an empty vector (EV) control into the contralateral muscle. TA muscles were harvested after 7 and 30 days.

### Mouse Skeletal Muscle Tissue Collection

Following completion of the appropriate time period, mice were anesthetized with isoflurane, and the TA muscles were excised, weighed, frozen in liquid nitrogen, and stored at −80°C for later analyses. Muscles were collected for histology (n = 5) and RNA/Protein isolation (n = 5) and processed as described below. On completion of tissue removal, mice were euthanized by exsanguination.

### Immunohistochemistry and Histology Tissue

Harvested mouse TA muscles were immediately weighed and fixed in 4% (w/v) paraformaldehyde for 16 h at 4°C and then placed in 30% sucrose for overnight incubation. The TA muscles were then embedded in Tissue Freezing Medium (Triangle Biomedical Sciences), and a Thermo HM525 cryostat was used to prepare 10 μm sections from the muscle mid-belly. All sections were examined and photographed using a Nikon Eclipse Ti automated inverted microscope equipped with NIS-Elements BR digital imaging software.

### Laminin Stain of Tissue

TA muscle sections were permeabilized in PBS with 1% triton for 10 minutes at room temperature. After washing with PBS, sections were blocked with 5% goat serum for 15 minutes at room temperature. Sections were incubated with Anti-Laminin (1:500, Sigma-Aldrich Cat# L9393, RRID: AB_477163) in 5% goat serum for 2 hours at room temperature, followed by two 5-minute washes with PBS. Goat-anti-rabbit AlexaFluor® 555 secondary (1:333, Invitrogen Cat# A28180, RRID: AB_2536164) in 5% goat serum was then added for 1 hour at room temperature. Slides were cover slipped using ProLong Gold Antifade reagent (Life Technologies). Image analysis was performed using Myovision software (41). Skeletal muscle fiber size was analyzed by measuring ≥ 250 transfected muscle fibers per muscle, per animal (10x magnification). Transfected muscle fibers were identified as GFP-positive as shown in results figures for 7 days and 30 days. We also measured the size of non-transfected fibers (GFP-negative) from the transfected muscles at all time points. Comparison of the CSA distributions of GFP-negative fibers in EV and UBR5 RNAi transfected muscles revealed no difference between groups. Therefore, fiber size comparisons were made between the GFP-positive fibers in the EV controls and UBR5 RNAi transected muscles.

### RNA Isolation and Total RNA Quantification of Tissue

Prior to RNA isolation, aliquots of frozen muscle powder were weighed in order to calculate RNA per milligram of wet muscle tissue. Muscle powder was homogenized using RNAzol RT reagent (Sigma-Aldrich, St Louis, MO) in accordance with the manufacturer’s instructions. Total RNA quantity and quality was assessed for 260/280 ratios using a SpectraMax M2 Microplate reader (Molecular Devices, CA, USA).

### Muscle Protein Synthesis (MPS) in Tissue

Changes in MPS were assessed in TA muscles transfected for 7 days by measuring the incorporation of exogenous puromycin into nascent peptides as described previously (19, 42). Puromycin (EMD Millipore, Billerica, MA, USA; cat. no. 540222) was dissolved in sterile saline and delivered (0.02 μmol g^−1^ body weight by I.P. injection) 30 min prior to muscle collection.

### Immunoblotting in Tissue Lysates

Frozen TA muscles were homogenized in sucrose lysis buffer (50 mM Tris pH 7.5, 250 mM sucrose, 1 mM EDTA, 1 mM EGTA, 1% Triton X 100, 50 mM NaF). The supernatant was collected following centrifugation at 8,000 *g* for 10 minutes and protein concentrations were determined using the 660-protein assay (Thermo Fisher Scientific, Waltham, MA). Twelve micrograms of protein were subjected to SDS-PAGE on 4-20% Criterion TGX stain-free gels (Bio-Rad, Hercules, CA) and transferred to polyvinylidene diflouride membranes (PVDF, Millipore, Burlington, MA). Membranes were blocked in 3% nonfat milk in Tris-buffered saline with 0.1% Tween-20 added for one hour and then probed with primary antibody (concentrations detailed below) overnight at 4°C. Membranes were washed and incubated with HRP-conjugated secondary antibodies at 1:10,000 (Mouse, Cell Signaling technology Cat# 7076, RRID AB_330924; Rabbit, Cell Signaling technology Cat# 7074, RRID AB_2099233) for one hour at room temperature. Immobilon Western Chemiluminescent HRP substrate was then applied to the membranes prior to image acquisition. Image acquisition and band quantification were performed using the Azure C400 System (Azure Biosystems, Dublin, CA, USA) and Image Lab, version 6.0.1 (Bio-Rad), respectively. Total protein loading of the membranes captured from images using stain-free gel technology was used as the normalization control for all blots. The following primary antibodies were used all used at a concentration of 1:1000: Cell Signaling Technologies (Danvers, MA, USA) – phospho-ERK1/2 ^Thr202/Tyr204^ (Cat# 4370, RRID:AB_2315112), ERK1/2 (Cat# 4695, RRID:AB_390779), phospho-p90RSK ^Ser380^ (Cat# 11989, RRID:AB_2687613), p90RSK (Cat# 9355, RRID:AB_659900), phospho-MSK1 ^Thr581^ (Cat# 9595, RRID:AB_2181783), phospho-MEK1/2 ^Ser217/221^ (Cat# 9154, RRID:AB_2138017), phospho-Akt ^Ser473^ (Cat# 4060, RRID:AB_2315049), Akt (Cat# 9272, RRID:AB_329827), phospho-GSK3β ^Ser9^ (cat. Cat# 9336, RRID:AB_331405), GSK3β (Cat# 9332, RRID:AB_2335664), phospho-p70S6K Thr^389^ (Cat# 9205, RRID:AB_330944), p70S6K (Cat# 9202, RRID:AB_331676), phospho-rpS6 ^Ser240/244^ (Cat# 5364, RRID:AB_10694233), rpS6 (Cat# 2217, RRID:AB_331355), phospho-4EBP1 ^Thr37/46^ (Cat# 2855, RRID: AB_560835), 4EBP1 (Cat# 9644, RRID:AB_2097841) UBR5 (Cat# 65344, RRID:AB_2799679), PP2Ac (Cat# 2259, RRID:AB_561239), eIF4e (Cat# 9742, RRID:AB_823488) and EMD Millipore – puromycin (Cat# MABE343, RRID:AB_2566826). Knockdown of UBR5 protein was confirmed previously with a reduction of 60% (Seaborne *et al.*, 2019) and all signaling antibodies in the present study were processed on the same samples as this UBR5 protein data.

### Statistical Analysis

Paired student *t-tests* were first carried out to assess knockdown efficiency (%) in siUBR5 treated vs. DM non-treated cells at in loaded and non-loaded myotubes. Also, to determine UBR5 mRNA fold-change in loaded vs. non-loaded myotubes dosed with DM alone. Finally, paired *t-tests* were carried out when comparing empty control (EV) vs. UBR5 RNAi transfected mouse TA SkM. All statistical analysis was performed using GraphPad Software (Prism, Version 7.0a, San Diego, CA). Data is presented as mean ± standard error of the mean (SEM). *P* ≤ 0.05 represents statistical significance.

## Results

### siRNA induced reductions in UBR5 gene expression are prevented with mechanical loading

We first wished to determine the impact of UBR5 silencing and mechanical loading on UBR5 gene expression in human myotubes (Figure 1A-F). There was an increase in mean UBR5 gene expression at 3 hrs post-loading in DM control conditions (158 ± 54.5% / ∼1.58-fold increase, Figure 1G), however this did not quite reach statistical significance. Following 3 hrs of UBR5 siRNA treatment (Figure 1H) in non-loaded myotubes, there was a significant decrease in UBR5 gene expression (−77 ± 1.28% *P* < 0.001; Figure 1G). Importantly, gene silencing of UBR5 at 3 hrs was prevented by mechanical loading, demonstrated by a return to baseline levels of UBR5 expression (Figure 1G) in the siUBR5 + loading condition (Figure 1G).

**Figure 1.**
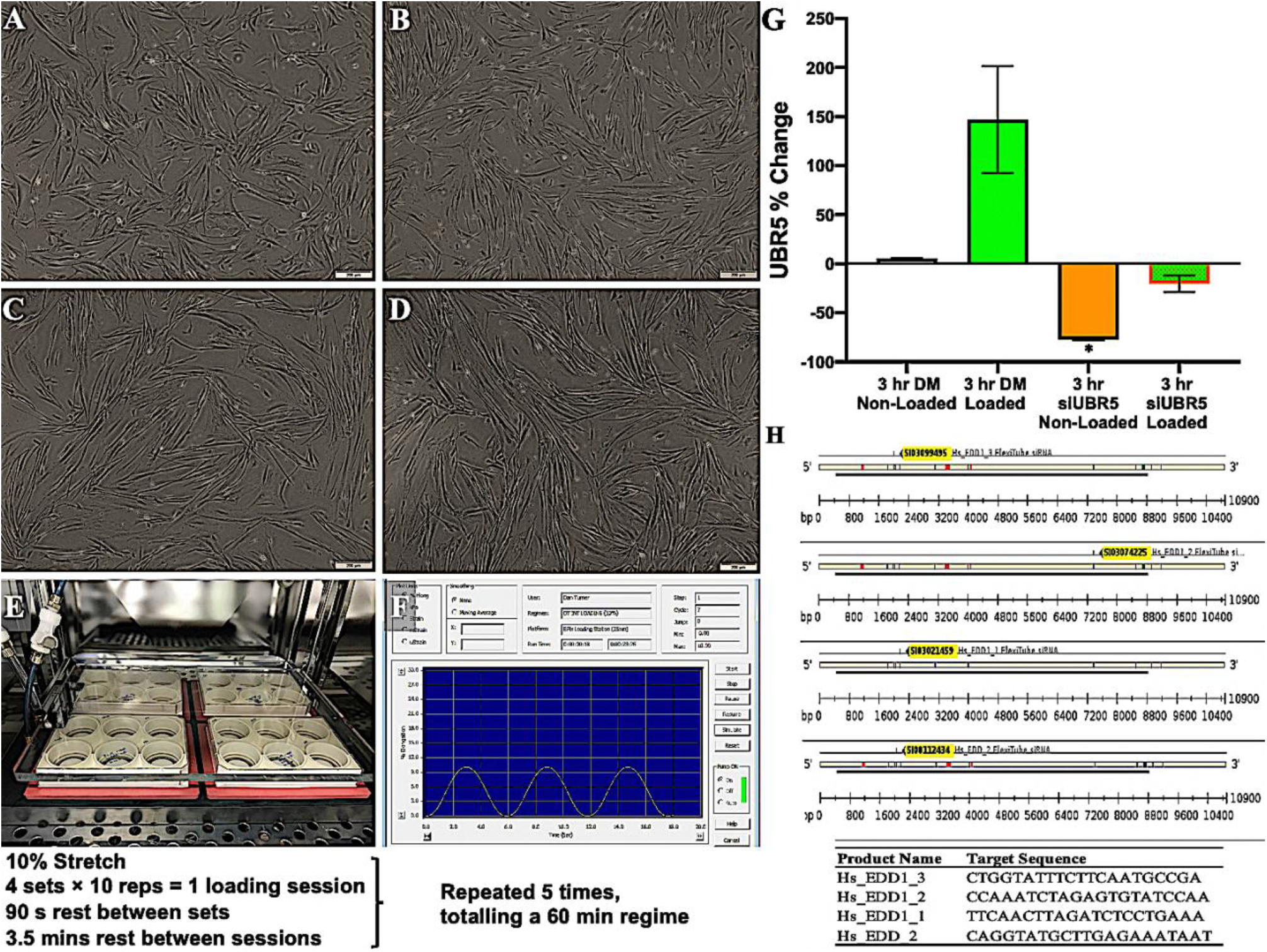
UBR5 gene silencing and mechanical loading of human myotubes. Cultured HMDC at **(A)** 0 hrs, **(B)** 72 hrs, **(C)** 7 days and **(D)** 10 days in DM (10× magnification, Olympus, CKX31) **(E)** HMDC cultured/differentiated into myotubes at day 10 and mechanically loaded on BioFlex^®^ well plates using Flexcell^®^ technology within a humidified incubator at 37°C/5 Co_2_%. **(F)** Example of the sine wave profile during the mechanical loading regime. **(G)** UBR5 gene expression increases (158 ± 54.6%) in loaded versus non-loaded myotubes. Silencing of UBR5 using siRNA reduced UBR5 mRNA expression (∼77%) in human myotubes after 3 hours. Mechanical loading prevented the reduction in UBR5 gene expression in UBR5 siRNA conditions back towards baseline levels at 3 hrs post-loading. **(H)** Graphical representation of the 4 × siRNA’s (siUBR5) used, and the sequence locations on the EDD1/UBR5 human gene (NM_015902, 10900 bp length). (**\***) Depicts statistical significance (*P* ≤ 0.05). Data is present as mean ± standard error of mean (SEM).

### Fiber CSA was significantly reduced at 7 days post electroporation in mice

To examine the effect of reducing UBR5 expression *in-vivo*, mouse TA muscles were transfected with either UBR5 RNAi plasmid targeting UBR5s HECT domain (Figure 2A) or an EV control. UBR5 protein was significantly reduced by 65% (± 11.8%) in UBR5 RNAi conditions (Figure 2B), as previously demonstrated in (35). The TA mass (mg) did not significantly change at 7 days post electroporation in UBR5 RNAi vs. EV transfected TA muscle (Figure 2C). Interestingly, however, measurement of the CSA of GFP-positive fibers revealed that the mean fiber CSA of GFP-positive fibers was significantly reduced (−9.5 ± 3.2%) in RNAi transfected TA muscle (*P* = 0.048; Figure 2D & E) with fewer larger fibers (≥3400 µm^2^) present versus the EV (Figure 2D & F).

**Figure 2.**
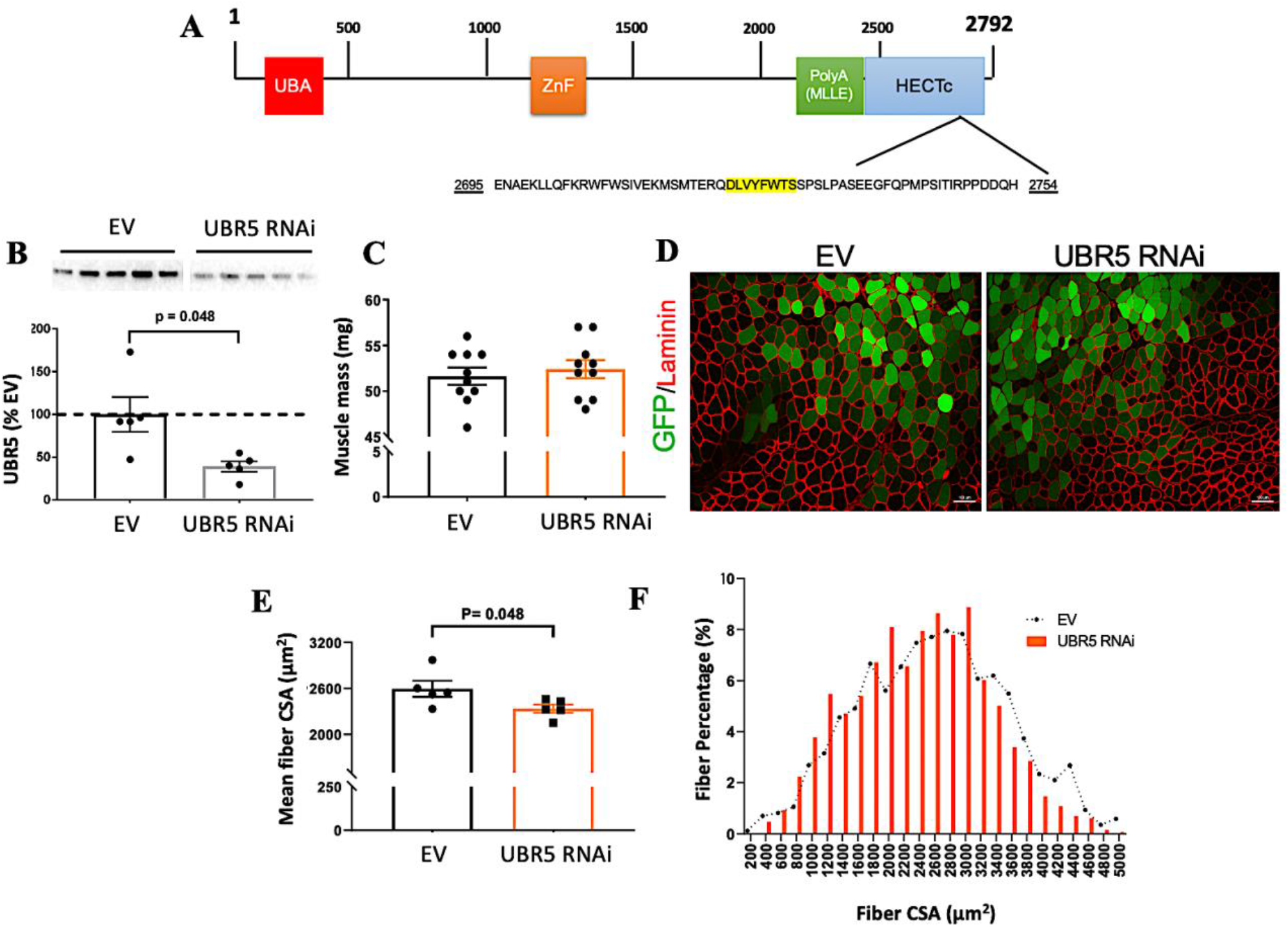
Characteristics of tibialis anterior muscle mass and fiber CSA in UBR5 RNAi transfected TA muscle after 7 days. **(A)** Schematic/ putative domain structure of UBR5 protein consisting of ubiquitin-associated (UBA) domain-like superfamily, Putative zinc finger in N-recognin (ZnF), Poly-adenylate binding protein (PolyA), MLLE protein/protein interaction domain and a HECT domain. The UBR5 RNAi plasmid was designed to target the nucleotide sequence in the HECT domain between amino acids position 2695-2754 (highlighted in yellow in A). **(B)** TA skeletal muscles were transfected for 7 days and we confirmed that the RNAi plasmid significantly reduced UBR5 protein levels by 65% (Seaborne et al., 2019) in basal mouse skeletal muscle. We have previously confirmed significant reductions in UBR5 protein levels at 7 days with this UBR5 RNAi plasmid (Seaborne et al., 2019), and all 7 day data in the present manuscript (muscle mass (C), histology (D), fiber CSA (E) RNA and signalling data – figure below) is analyzed from the same animals/samples as this UBR5 protein data. Figure 2B is therefore reused with permissions from; R. A. Seaborne et al., *Journal of Physiology* (Wiley), 597.14 (2019) pp 3727–3749, Copyright-2019 The Authors. *The Journal of Physiology*. Copyright-2019 The Physiological Society). **(C)** After 7 days, muscle mass was no different between empty vector control (EV) and UBR5 RNAi transfected TA muscles (*n =* 10 per group). **(D)** Representative images (10x magnification; scale bar = 100 μm) for GFP transfected fiber identification and CSA quantification through Laminin staining. **(E)** A significant reduction (*P* = 0.05) in mean transfected fiber CSA size was observed in the RNAi transfected vs. EV muscles *(n =* 5 per group). **(F)** Quantification of muscle CSA revealed changes in the percentage of small fibers (<1200 μm^2^) and large fibers (≥3400 μm^2^) with RNAi transfected muscles verses EV group (N = 5 per group). Data presented as Mean ± SEM. Statistical significance is depicted where present (P ≥ 0.05).

### Total RNA concentrations, muscle protein synthesis, ERK1/2 and Akt activity and total p90RSK are reduced in UBR5 RNAi TA muscle after 7 days

Along with a reduction in fiber CSA at 7 days post UBR5 RNAi, a significant reduction in total RNA concentrations (*P* = 0.01, Figure 3A) and protein synthesis (−17 ± 3.2%, *P* = 0.05, Figure 3B & 3C) was evident. Wishing to determine potential regulators of this process in the ERK signalling pathway (Figure 3D), ERK1/2 (p44/42 MAPK) was examined and found to be reduced (−43 ± 2.3%, *P =* 0.03, Figure 3E). Interestingly, the reduction in RNA, protein synthesis and ERK1/2 signalling coincided with a significant increase in phospho-p90RSK^Ser380^ (170 ± 39.8%, *P* = 0.04; Figure 3F) at 7 days. However, this was on the background of a significant 45% decrease in total p90RSK (−45 ± 5.6%, P = 0.02; Figure 3G) with no significant change in total ERK1/2 protein levels were observed in UBR5 RNAi transfected muscles. We also observed no differences in the phosphorylation of MEK 1/2^Ser 217/221^ or MSK-1^Thr581^ between RNAi and EV transfected TA muscles after 7 days (data not depicted). We next assessed protein signalling associated with the Akt pathway (Figure 4A). As with ERK1/2, there were significant reductions in Akt ^ser473^ phosphorylation in UBR5 RNAi transfected TA muscle (−26 ± 9%, P = 0.03; Figure 4B), with no differences in GSK-3β ^Ser9^ phosphorylation after 7 days post-electroporation (Figure 4A). Interestingly, despite reductions in ERK1/2, Akt and protein synthesis, we observed significant elevations in phosphorylation for p70S6K ^Thr389^ (176 ± 57%, P = 0.02; Figure 4C) and rpS6 ^Ser240/244^ (178 ± 39.4%, P = 0.01; Figure 4D) in UBR5 RNAi transfected muscles at 7 days. Finally, there were no changes in activity of 4E-BP1 ^Thr37/46^, eiF4e or PP2Ac at 7 days in UBR5 RNAi transfected muscles (Figure 4A).

**Figure 3.**
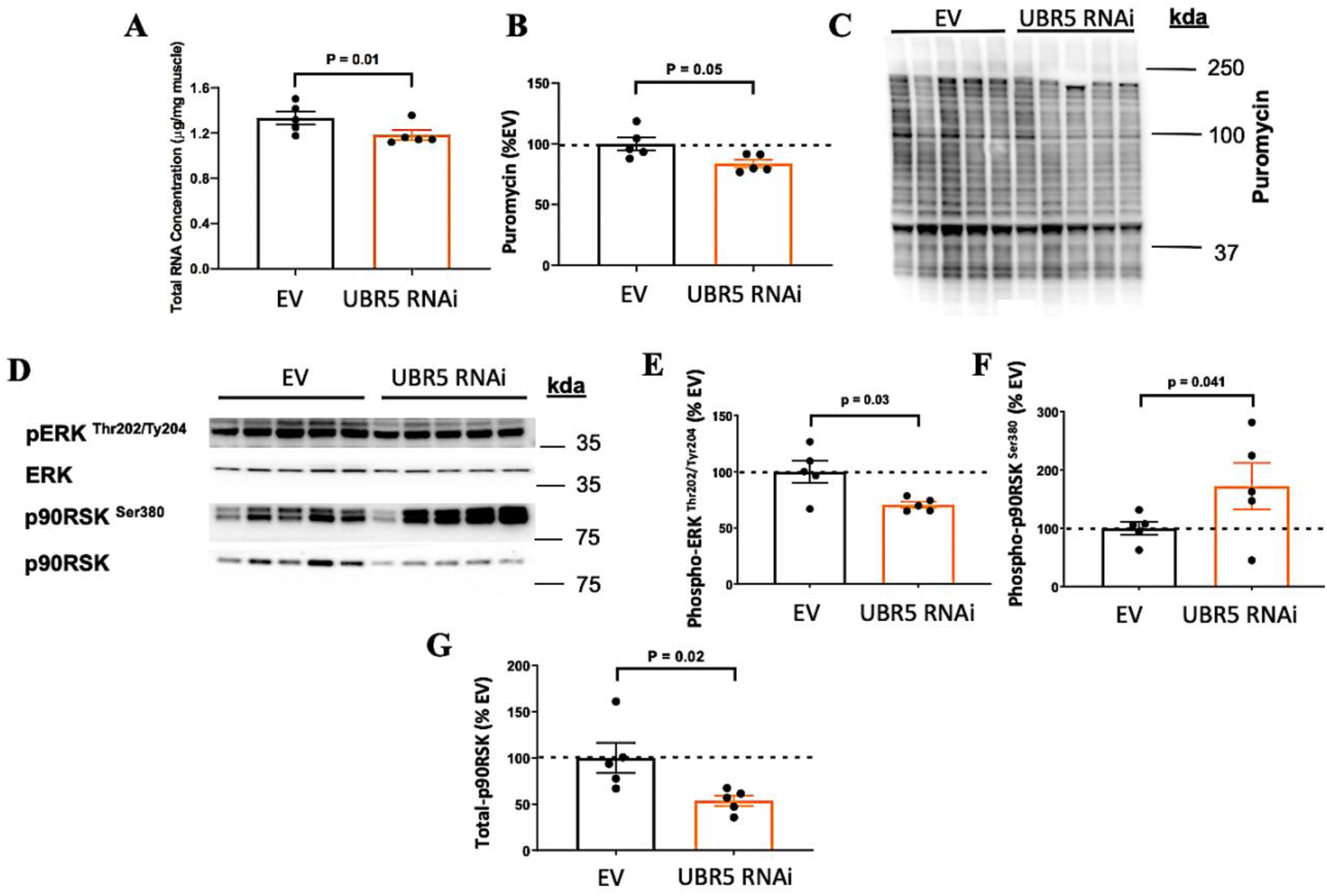
Alteration in total RNA, protein synthesis and MAPK (ERK/p90RSK) signaling in UBR5 RNAi transfected TA muscle after 7 days. **(A)** Total RNA concentration (assessed as µg/mg of muscle) was significantly reduced (*P* = 0.01) in the UBR5 RNAi muscles. **(B & C)** RNAi transfected muscles also displayed a significant reduction (∼17%, *P* = 0.05) in muscle protein synthesis (B) assessed via western blots for puromycin incorporation (C). **(D)** Western blot images for the ERK signaling pathway. **(E)** Decreased phosphorylation levels of ERK1/2 (p44/42 MAPK) *(P =* 0.03) and **(F)** increased phosphorylation levels of p90RSK (*P* = 0.04) with UBR5 RNAi. **(G)** Reduced levels of total p90RSK (*P* = 0.02). Data presented as Mean ± SEM. Total protein loading of the membranes captured from images using stain-free gel technology was used as the normalization control for all blots. N = 5 per group. Statistical significance is depicted where present (P ≥ 0.05).

**Figure 4.**
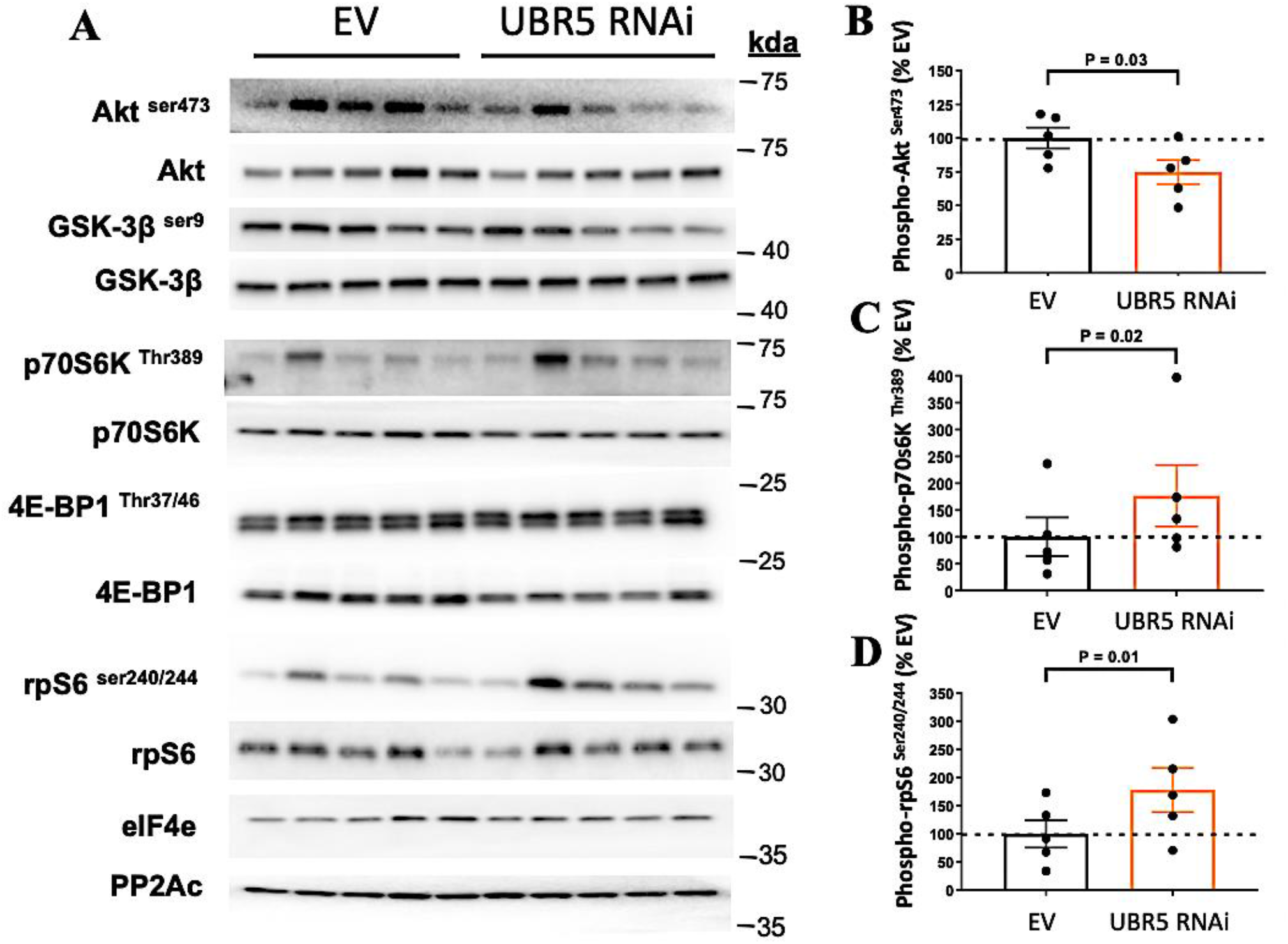
(A) Western blot analysis for the Akt signaling pathway (Akt, GSK-3β, p70S6K, 4E-BP1, rpS6, eIF4e, PP2Ac) in UBR5 RNAi transfected TA muscle after 7 days. **(B)** Decreased phosphorylation levels of Akt (*P* = 0.03) and **(C)** increased phosphorylation levels of p70S6K and **(D)** rps6 with UBR5 RNAi. No significant changes in phosphorylation of GSK-3β, 4E-BP1 or total levels of eiF4e and PP2Ac were observed in UBR5 RNAi conditions at 7 days. All *n* = 5 per group – EV control and UBR5 RNAi. Data presented as Mean ± SEM. Total protein loading of the membranes captured from images using stain-free gel technology was used as the normalization control for all blots. Statistical significance is depicted where present (P ≥ 0.05).

### Prolonged UBR5 RNAi transfection leads to significant loss of muscle mass and fiber CSA atrophy in mouse skeletal muscle

Given the data over 7 days, the next question to interrogate was the impact of UBR5 RNAi transfection in TA muscle after a prolonged period (30 days), with the hypothesis that this would perhaps lead to greater muscle atrophy vs. 7 days of UBR5 suppression. In line with this hypothesis, while UBR5 protein reductions were not maintained out to this 30-day timepoint (Figure 5A), the earlier reductions in UBR5 at 7 days led to a significant reduction in muscle mass by 30 days (−4.6 ± 1.5%) in UBR5 RNAi transfected vs. EV muscles (P = 0.01; Figure 5B). Alongside the reduction in muscle mass, a significant larger reduction in GFP-positive fiber CSA (−18.2 ± 2.3% at 30 d vs. -9.5 ± 3.2% at 7 d) was evident after UBR5 RNAi transfection (*P* = 0.01; Figure 5C & D) vs. EV control, with the RNAi transfected muscles displaying a shift in the distribution of fiber CSA primarily towards smaller fibers (≥ 2600 µm^2^) compared to the EV control muscle (Figure 5E). These observations provide further support towards UBR5 being a positive modulator of muscle mass, with its knockdown evoking considerable atrophy.

**Figure 5.**
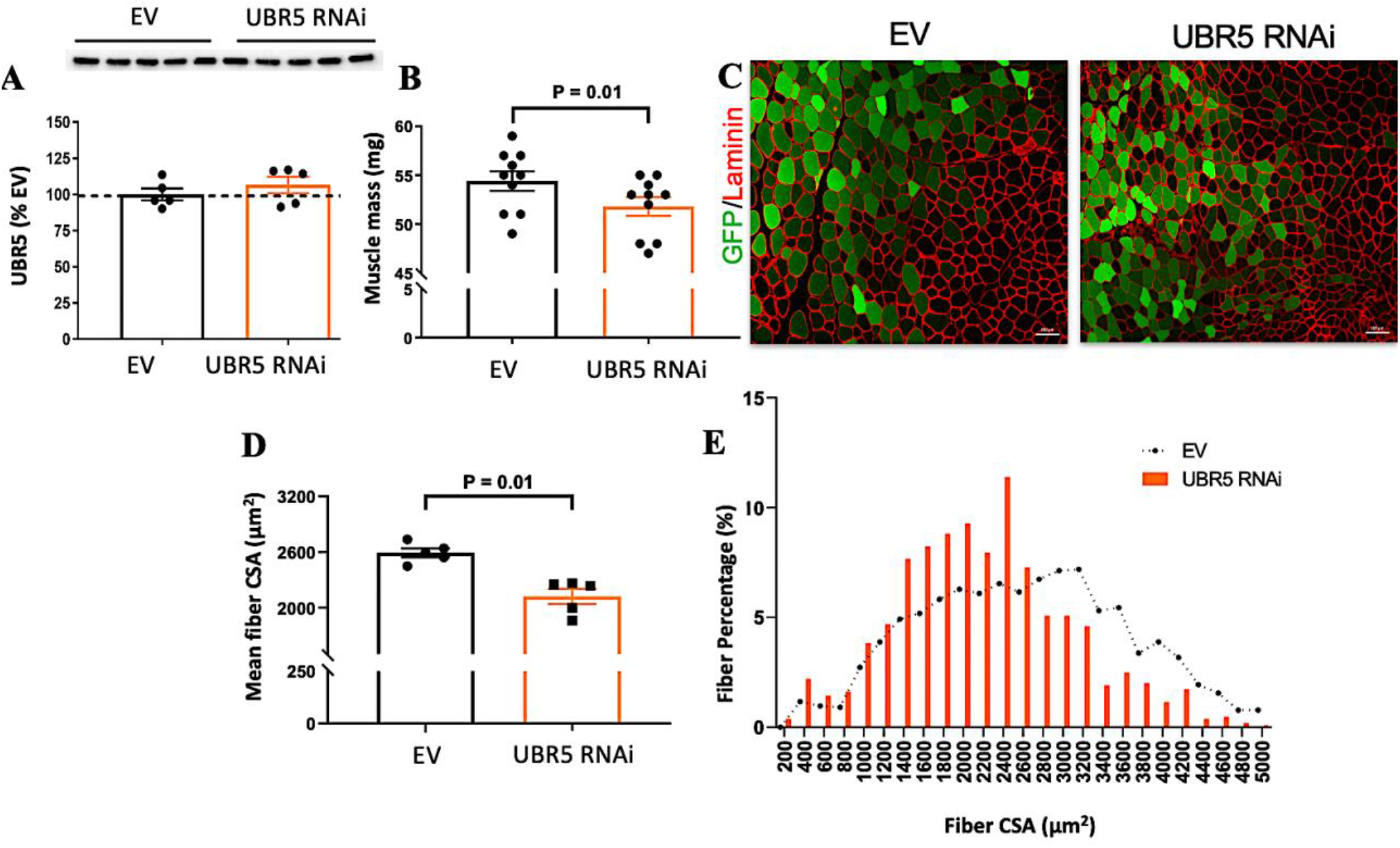
Loss of muscle mass and fiber CSA in UBR5 RNAi transfected TA muscle after 30 days. **(A)** UBR5 protein after 30 d UBR5 RNAi. (**B)** After 30 days, muscle mass was significantly reduced in UBR5 RNAi transfected vs. empty vector control (EV) TA muscles (P = 0.01; *n =* 10 per group). **(C)** Representative images (10x magnification; scale bar = 100 μm) for GFP transfected fiber identification and CSA quantification through Laminin staining. **(D)** A significant reduction (*P* = 0.01) in mean transfected fiber CSA size was observed in the RNAi transfected vs. EV muscles *(n =* 5 per group). **(E)** Quantification of transfected muscle CSA revealed a leftward shift from large to smaller fiber sizes (≥3400 μm^2^) with RNAi transfected muscles verses EV group (**D**). Data presented as Mean ± SEM.

### Prolonged UBR5 RNAi at 30 days and reductions in muscle mass and fiber size are associated with increased PP2Ac, reduced eIF4e and inappropriate chronic elevation of p70S6K and rpS6

At 30 days post RNAi transfection we continued to observe an elevation in p90RSK phosphorylation, similar to the 7d time point (Figure 6A). Opposite to the 7d time point, we observe significant increases total p90RSK (Figure 6B) and both phosphorylated and total ERK (Figure 6C & 6D) in RNAi transfected muscles after 30 days. As with 7 days we observed no differences in the phosphorylation of MEK 1/2^Ser 217/221^ or MSK-1^Thr581^ between RNAi and EV transfected TA muscles after 30 days (data not depicted). By 30 days there was no reduction of phosphorylated Akt as observed at the 7-day timepoint and no change in GSK-3β phosphorylation with RNAi transfection (Figure 7A). Interestingly, elevation of p70S6K and rpS6 phosphorylation from the 7-day time point continued over the more chronic period of 30 days, demonstrating a 4- and 2-fold increase respectively in the RNAi transfected muscles verses the empty vector control group (Figure 7B & 7C). Compared to the 7d time point, we also observed an 80% reduction in eIF4e protein levels in the RNAi group (Figure 7D). Furthermore, the levels of PP2Ac were significantly increased at the 30 d time point (Figure 7E). Finally, like at 7 days there were no changes in activity of 4E-BP1 ^Thr37/46^ or its total levels in UBR5 RNAi transfected muscles at 30 days. Signalling events at 7 and 30 days are summarised in cell signalling diagram (Figure 8).

**Figure 6.**
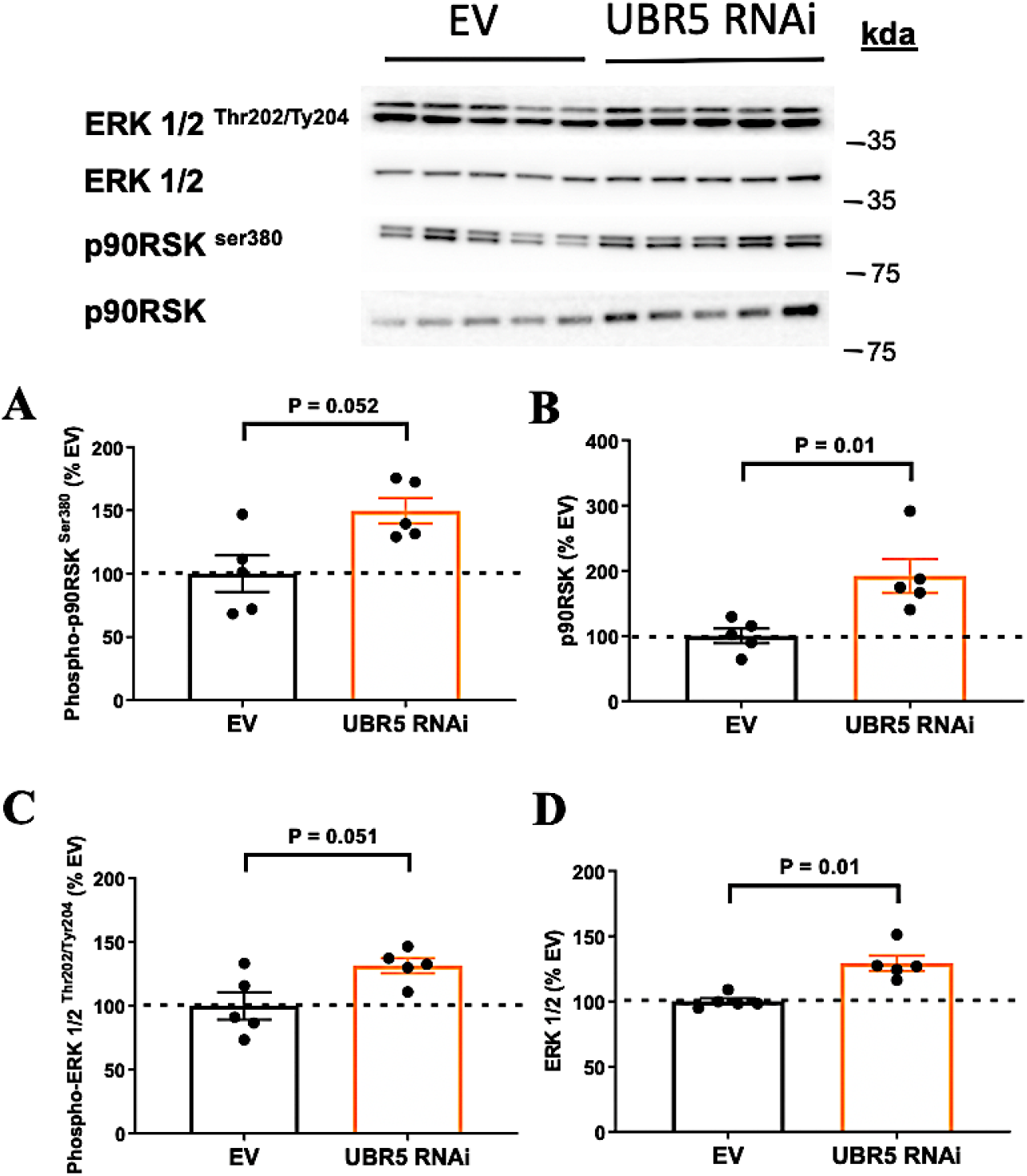
ERK pathways signaling in UBR5 RNAi transfected TA muscle after 30 days. **(A)** Phosphorylated p90RSK, **(B)** total p90RSK, **(C)** Phosphorylated ERK1/2 and **(D)** Total ERK. All *n* = 5 per group – EV control and UBR5 RNAi. Data presented as Mean ± SEM. Total protein loading of the membranes captured from images using stain-free gel technology was used as the normalization control for all blots. Statistical significance is depicted where present (P ≥ 0.05).

**Figure 7.**
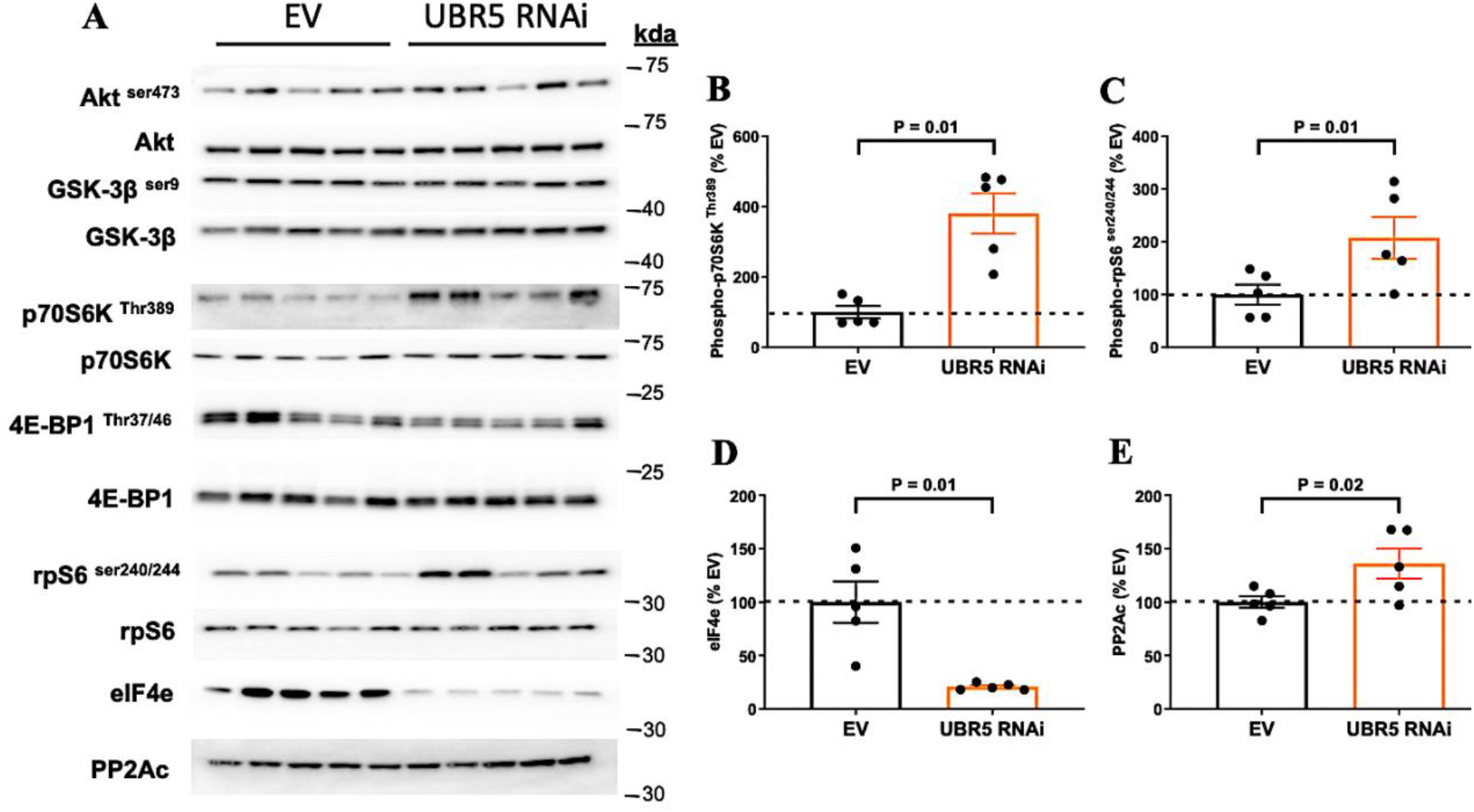
A) Western blot analysis for the Akt signaling pathway (Akt, GSK-3β, p70S6K, 4E-BP1, rpS6, eIF4e, PP2Ac) in UBR5 RNAi transfected TA muscle after 30 days. Significant changes observed for **(B)** Phosphorylated p70S6K, **(C)** Phosphorylated rpS6, **(D)** eIF4e and, **(E)** PP2Ac. All *n* = 5 per group – EV control and UBR5 RNAi. Data presented as Mean ± SEM. Total protein loading of the membranes captured from images using stain-free gel technology was used as the normalization control for all blots. Statistical significance is depicted where present (P ≥ 0.05).

**Figure 8.**
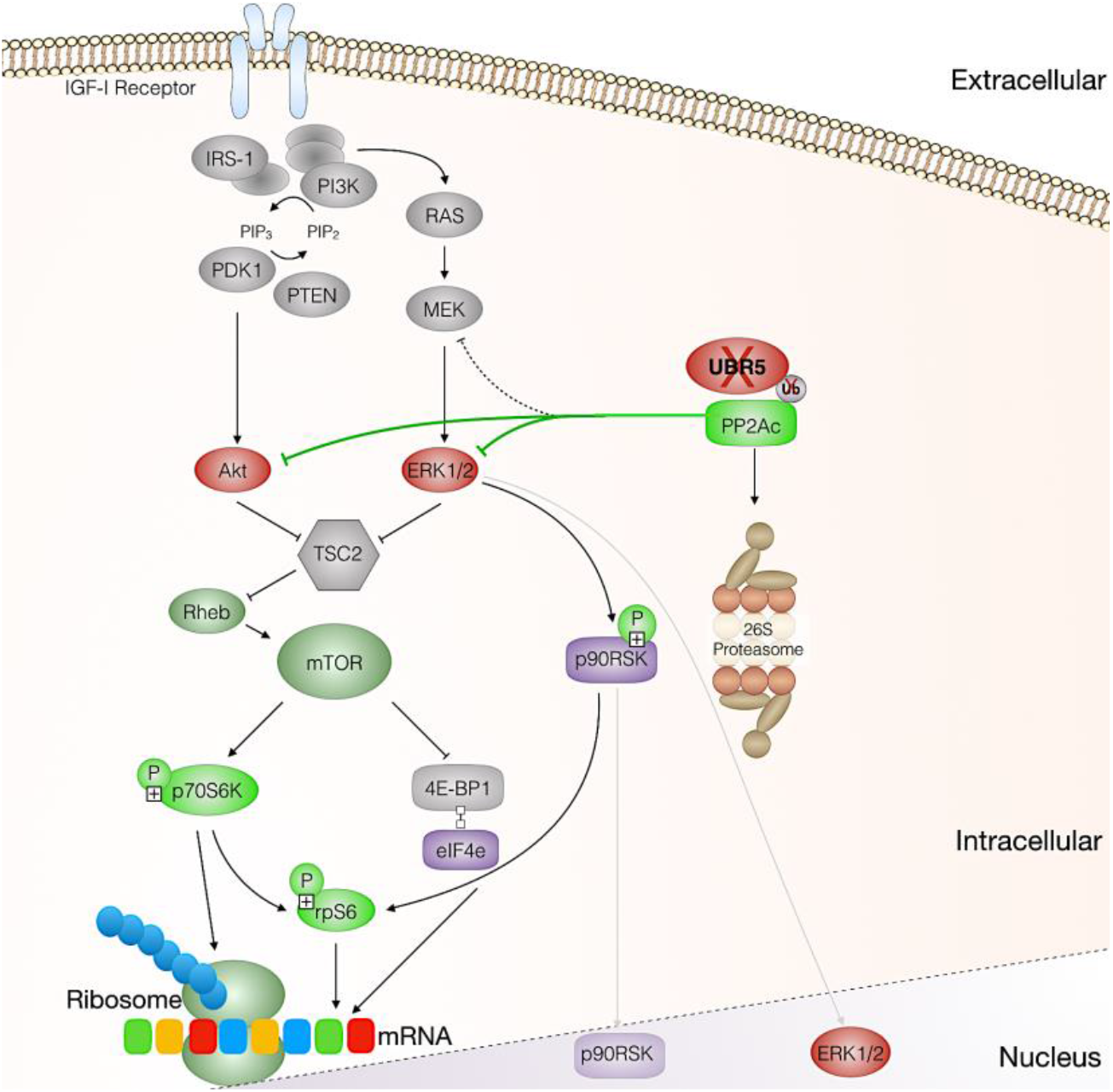
Schematic of cell signaling events after UBR5 knockdown in skeletal muscle in-vivo. UBR5 knockdown (X) increases phosphatase PP2Ac, reduces ERK and Akt phosphorylation (RED), reduces total p90RSK and eIF4e (PURPLE), chronically increases the phosphorylation of p70S6K and rpS6 (GREEN).

## Discussion

The present study aimed to: 1) silence UBR5 in human myotubes using siRNA during mechanical loading to confirm whether UBR5 was regulated in response to mechanical loading. Given ERK/p90RSK and Akt/P70S6K signalling has been associated with load-induced anabolism, ERK/p90RSK is altered with UBR5 manipulation in non-muscle cells, and PP2Ac is a negative regulator of ERK and Akt that can be degraded by UBR5, we also aimed to: 2) investigate the impact of reduced UBR5 on ERK and Akt signalling pathways in SkM tissue using our miR-based RNAi for UBR5 electroporated into the tibialis anterior (TA) of mice.

We hereby report that mechanical loading of non-treated myotubes induced anincrease in UBR5 gene expression which was similar to that following acute loading in murine bioengineered SkM (35) and acute RE in humans (36, 37). Furthermore, mechanical loading was able to prevent the siRNA-induced reduction in UBR5 gene expression, which suggested that UBR5 gene expression may be regulated via mechano-sensitive signalling. Indeed, transfection of UBR5 RNAi into the TA muscle of mice after 7 days caused a significant reduction in fiber CSA, total RNA, global protein synthesis and ERK1/2 / Akt phosphorylation. However, p90RSK, P70S6K and rps6 phosphorylation significantly increased, perhaps suggestive of a potential compensatory mechanism triggered by the reduction in UBR5 or inappropriate chronic activation of these pathways, like that observed in aged muscle (discussed below). Furthermore, the increases in p90RSK phosphorylation were on a background of considerably reduced total p90RSK levels. Finally, changes at 7 days also culminated in significant reductions in muscle mass and an even greater reduction in fiber CSA after prolonged (30 days) UBR5 RNAi. This was associated with increased PP2Ac and, as hypothesised above, continued chronic elevation of P70S6K and rpS6 from 7 to 30 days, as well as reductions in elongation initiation factor 4e (eIF4e). Overall, the present study further supports the notion that UBR5 plays an important role in muscle anabolism and hypertrophy, and its reduction culminates in atrophy.

In the present study, transfection of UBR5 targeting siRNAs induced a ∼77% reduction in UBR5 mRNA expression in non-loaded human myotubes. Following confirmation of effective UBR5 knockdown in non-loaded myotubes, UBR5 gene expression was assessed in response to mechanical loading. Mechanical loading of non-treated myotubes displayed an increase (∼158% / 1.58-fold) in UBR5 gene expression (see Figure 1G), which was similar to the fold changes observed after an acute bout of RE in human SkM (∼1.71-fold) (Seaborne *et al.*, 2018*b*, 2018*a*) and the same fold changes observed after acute loading in bioengineered SkM (∼1.58-fold) (35). Significant changes in UBR5 gene expression were also achieved following chronic (7 weeks) training, and in retraining for which greatest increase in UBR5 gene expression was observed (36, 37). Also, UBR5 significantly increased after hypertrophy following 4 weeks of resistance training via programmed chronic high-frequency electrical stimulation in rats (34, 35). Furthermore, we have previously reported a significant increase in UBR5 following acute loading in bioengineered C_2_C_12_ SkM (35). Finally, mechanical loading was able to prevent the siRNA-induced reduction in UBR5 gene expression. Such a finding perhaps suggested that UBR5 may be involved in mechano-sensitive signalling pathways.

Therefore, we next performed UBR5-specific knockdown experiments in mouse TA muscle, to investigate the impact of reduced UBR5 on mechanical-induced signalling networks, ERK (MEK/ERK/p90RSK/MSK1) and Akt (PP2Ac/Akt/GSK3β/P70S6K/4E-BP1/rpS6) pathways *in-vivo*. Electroporation of UBR5 RNAi for 7 days induced a significant reduction in average myofiber CSA, frequency of larger myofibers (≥3400 µm^2^), total RNA and muscle protein synthesis (∼17%) which coincided with a reduction in ERK1/2 signalling. The reduction in total RNA and muscle protein synthesis is interesting given the classification of UBR5 as an E3 ubiquitin ligase and thus its role in the ubiquitin-proteasome system. Indeed, UBR5 may have unidentified functions within skeletal muscle that are unrelated to proteasomal degradation. An example of an E3 ubiquitin ligase having other functions in skeletal muscle is TRIM72 (known as MG53), with its role being studied in cell membrane repair (6, 11, 12). UBR5 has been suggested to have a role in the regulation of mRNA translation and gene regulation through the MLLE domain on the UBR5 protein structure (30, 44). Recent studies have identified the translation capacity and activity of skeletal muscle to be important for growth and hypertrophy in rodent and human models (20, 31, 39, 42). Therefore, while we saw reductions in ERK/ Akt activity in the present study, there were opposite increases in p90RSK, p70S6K and rps6 suggesting that the observed reductions in protein synthesis may not be regulated by p90RSK, p70S6K and rps6 activity. However, there were significant reductions in total p90RSK as a consequence of UBR5 knockdown at 7d, that may contribute to the reductions in protein synthesis observed. However, given the observed reduction in total RNA, there could be a role for UBR5 in the regulation of mRNA translational capacity that would in-turn lead to reductions in total protein synthesis and SkM mass. Indeed, supporting this notion, we also observed 80% reductions of eiF4e at 30 days, an elongation initiation factor important for enhanced RNA translational efficiency, splicing and stability.

Other explanations for increased p90RSK, p70S6K and rpS6 signalling activity significantly increasing following UBR5 knockdown, are perhaps due to a compensatory or protective mechanism in an attempt to prevent the reductions in protein synthesis. Further, in a non-classically envisaged pathway, the increase in p90RSK activity may be explained due to its suggested role as an upstream regulator of UBR5 and its activation could therefore be altered due to the suppression of UBR5 protein levels (9). This could suggest that loss of UBR5 may indirectly lead to activation of p90RSK because its normal target for action is no longer present. Alongside the alterations in p90RSK phosphorylation, as suggested above, we did observe a significant loss in total protein levels for p90RSK at 7 days which may contribute towards the alterations in p90RSK activity and the overall reductions in protein synthesis. There are currently no commercial antibodies to assess UBR5 activity in relation to p90RSK, something that warrants investigation when antibodies are developed in the future. In addition to alterations in ERK and Akt signalling and reduced total p90RSK and protein synthesis at 7 days, a prolonged time period after transfection of the UBR5 RNAi at 30 days resulted in greater reductions in both muscle mass and fiber CSA. The loss of muscle mass and fiber size at 30 days, was surprisingly accompanied by increased p70S6K and rpS6 phosphorylation. The chronic activation of mTORC/P70S6K signalling has been observed to be detrimental in skeletal muscle homeostasis, such as that with increasing age (2, 8, 24, 38). Therefore, subsequent future studies will seek to address the interplay between UBR5 and Akt/mTORC/p70S6K signalling with aging.

Finally, in the present study we observed significant increases in PP2Ac levels at 30 d RNAi. PP2Ac is a phosphatase demonstrated to inhibit Akt and ERK signalling, and importantly, UBR5 in its capacity as an ubiquitin ligase has been demonstrated to target PP2A for degradation in non-muscle studies (25–27, 38). While observing an increase in PP2Ac at 30 days, the reductions in signalling of Akt and ERK were observed at 7 days and not at 30 days. However, the data does suggest for the first time in skeletal muscle, that PP2A is perhaps a target of UBR5 and that knockdown of UBR5 leads to an increase in the levels of PP2A due to a potential decrease in its degradation. However, UBR5 and its protein targets remain to be elucidated in skeletal muscle and require further study given these interesting findings.

Despite the exciting and novel findings surrounding UBR5’s mechanistic role in skeletal muscle, it is worth acknowledging the limitations of the study. Firstly, neither puromycin incorporation nor subsequent measures of total protein synthesis were assessed in human myotubes. This was due to the limited human cellular material available during experimentation. It was therefore not feasible to extract protein for all conditions, nor to examine cellular signalling in siUBR5 treated vs. non-treated human myotubes. However, we subsequently performed this signalling analysis *in-vivo*. It is also important to note that UBR5 protein levels were not reduced at 30 d to the same degree as the 7d time point and this may be reflective of the effectiveness of the RNAi construct being reduced and also the influence of non-transfected muscle fibers in the biochemical analyses. However, the earlier reductions in UBR5 at 7 days still evoked altered signalling and greatest loss of muscle mass by this 30-day timepoint. Suggesting the earlier reductions in UBR5 at 7 days still culminated in later atrophy by 30 days. Lastly, it would also be prudent in future experiments to also evoke hypertrophy (e.g. via synergistic ablation) in rodents in the presence or absence of the UBR5 RNAi plasmid.

## Conclusion

The present study supports the notion that UBR5 plays an important role in muscle anabolism/hypertrophy, and presents novel findings demonstrating knockdown of UBR5 is associated with early reductions in ERK1/2 and Akt activity, total p90RSK and protein synthesis that culminates in later atrophy that is associated with increased PP2Ac, reduced eIF4e levels and inappropriately chronically elevated activity of p70S6K and rpS6.

## Author contributions

DCT, DCH, LMB, SCB, APS conceived and designed the research. All authors were involved in acquisition or analysis or interpretation of data for the work. DCT, DCH, LMB, SCB, APS drafted the work. All Authors were involved in revising the work critically for important intellectual content. All authors approved the final version of the manuscript.

## Funding

DCT was funded via PhD studentships from Keele University and Liverpool John Moores University (LJMU) via APS. DCTs project was further supported by the Society for Endocrinology equipment grant and the North Staffordshire Medical Institute awarded to APS. RAS was funded by Doctoral Alliance/ LJMU funded PhD studentship via APS and further supported by grants awarded to APS from GlaxoSmithKline. PG was funded by EPSRC/MRC (UKRI) PhD studentship via APS and Keele University doctoral training centre and is now supported by the Norwegian School of Sport Sciences. DCH & LMB was supported by the University of Iowa in the laboratory of SCB.

## Declarations & Competing Interests

SCB is on the scientific advisory board for Emmyon Inc. The authors declare that they have no other competing interests.

## Data Availability Statement

The data that support the findings of this study are available from the corresponding author upon reasonable request.

## Acknowledgments

We wish to thank Dr. David S. Waddell (University of North Florida) and Mr. George R. Marcotte (University of Iowa) for technical assistance with the project.

## Notes

### Competing Interest Statement

Prof. Sue C Bodine is on the scientific advisory board for Emmyon Inc. The authors declare that they have no other competing interests.

### Summary of Updates

We have now analysed additional signalling molecules in the Akt pathway and added a signalling diagram.

